# Activation of macrophages by lysophosphatidic acid through the lysophosphatidic acid receptor 1 as a novel mechanism in multiple sclerosis pathogenesis

**DOI:** 10.1101/870980

**Authors:** Jennifer Fransson, Ana Isabel Gómez, Jesús Romero-Imbroda, Oscar Fernández, Laura Leyva, Fernando Rodriguez de Fonseca, Jerold Chun, Celine Louapre, Anne Baron Van-Evercooren, Violetta Zujovic, Guillermo Estivill-Torrús, Beatriz García-Díaz

**Affiliations:** Institut du Cerveau et de la Moelle Epinière-Groupe Hospitalier Pitié-Salpêtrière, INSERM, U1127, CNRS, UMR 7225, F-75013 Paris; Sorbonne Universités, Université Pierre et Marie Curie Paris 06, UM-75, F-75005 Paris, France; Unidad de Gestión Clínica de Neurociencias, Instituto de Investigación Biomédica de Málaga-IBIMA, Hospital Regional Universitario de Málaga, Málaga, Spain; Unidad de Gestión Clínica de Salud Mental, Instituto de Investigación Biomédica de Málaga-IBIMA, Hospital Regional Universitario de Málaga, Málaga, Spain; Sanford Burnham Prebys Medical Discovery Institute, La Jolla, CA, USA; Assistance Publique-Hôpitaux de Paris, Neurol. Dept. Pitié Salpétrière Univ. Hosp., Paris, France

## Abstract

Multiple sclerosis (MS) is a neuro-inflammatory disease for which the pathogenesis remains largely unclear. Lysophosphatidic acid (LPA) is an endogenous phospholipid that is involved in multiple immune cell functions and is dysregulated in MS. Its receptor LPA_1_ is expressed in macrophages and regulates their activation, which is of interest due to the role of macrophage activation in MS in both destruction and repair.

In this study, we studied the viable Malaga variant of LPA_1_-null mutation as well as pharmaceutical inhibition of LPA_1_ in mice with experimental autoimmune encephalomyelitis (EAE), a model of MS. LPA_1_ expression was also analyzed in both wild-type EAE mice and MS patient immune cells. The effect of LPA and LPA_1_ on macrophage activation was studied in human monocyte-derived macrophages.

We show that lack of LPA_1_ activity induces a milder clinical course in EAE, and that *Lpar1* expression in peripheral blood mononuclear cells (PBMCs) correlates with onset of relapses and severity in wild-type EAE mice. We see the same over-expression in PBMCs from MS patients during relapse compared to progressive forms of the disease, and in monocyte-derived macrophages after exposure to pro-inflammatory stimuli. In addition, LPA induced a pro-inflammatory-like response in macrophages through LPA_1_, providing a plausible way in which LPA and LPA_1_ dysregulation can lead to the inflammation seen in MS.

These data show a new mechanism of LPA signaling in the pathogenesis of MS, prompting further research into its use as a therapeutic target biomarker.

## Introduction

Multiple sclerosis (MS) stands out as one of the most widespread neurological diseases in young adults affecting approximately 2.3 million people worldwide (40). MS pathogenesis consists of inflammation of the central nervous system (CNS), oligodendrocyte death and myelin damage, and many different immune cells play an important role. Although T cells have always been considered central in MS pathogenesis, new insights have unveiled the key importance of macrophages in this disease. Macrophages can play a dual role in MS pathology; they can contribute to tissue damage and inflammation, but also exert a neuroprotective and regenerative effect (30). Their pleiotropic mode of action relies on their capacity to endorse different status of activation: classically activated, or M1, macrophages showing pro-inflammatory characteristics; and alternatively activated, or M2, macrophages displaying a more anti-inflammatory phenotype (46). Each activation state plays a different role along the process of remyelination. “M1” macrophages will be the first actor of the initiation of myelin repair: phagocytizing myelin debris and inducing oligodendrocytes precursor cells (OPC) proliferation and migration to the lesion site. Next, a switch in the macrophage population from the M1 to the M2 phenotype induces the secretion of trophic factors that foster OPC differentiation into new myelin-forming oligodendrocytes (31). However, to what extent macrophages intrinsically contribute to myelin destruction and repair in MS remains unclear.

Lysophosphatidic acid (LPA) is a bioactive phospholipid that influences numerous cell responses, including cell motility, neuropathic pain, inflammation, fibrosis, and cancer (20, 29, 44). The versatility of the LPA responses relies on its broad and dynamic presence in different tissues and fluids (2), as well as on its multiple receptors, both membrane (named from LPA_1_ to LPA_6_) (48) and nuclear (the peroxisome proliferator-activated receptor γ, PPAR-γ) (27). The expression of the LPA receptors being modular (37, 41), contributes to the adaptability of the LPA signaling.

In the past years, the role of LPA in the pathogenesis of different immune-related diseases has been recognized. LPA dysregulation has been implicated in different inflammatory diseases, such as atherosclerosis (8), cancer (45), and MS (41) because of its effect on immune cells, of both the innate and adaptive immune systems (3, 15, 23). However, despite the growing knowledge in this field, many aspects of the role of LPA in immune-related pathogenesis remain unclear. Interestingly, LPA_1_ – the first receptor of LPA to be discovered (18) – has a notable importance in the physiology and pathology of the CNS (49). While the importance of LPA in MS pathogenesis has not been neglected (4, 21, 41), showing a dysregulation of the serum LPA levels in MS patients (4, 41), with increased levels at relapse-onset and a role of LPA_2_ in T cells homing (41), the role of LPA_1_ remains unclear.

LPA has been shown to induce various effects in macrophages, and it constitutes a major serum survival factor for murine macrophages (22). Moreover, Lee et al. (23) showed that stimulation of mouse macrophages with LPA upregulated their expression of pro-inflammatory cytokines such as IL-1β and TNF-α transcripts and protein, and downregulated transcription of the anti-inflammatory IL-2. Interestingly, peripheral blood monocytes and/or tissue macrophages in both mice and humans express LPA_1_ receptor (1, 19). This receptor induces their activation, migration and infiltration in different disease mouse models (32, 42). The dysregulation of LPA and the importance of macrophage activation in MS thus present LPA_1_ as a potential receptor of interest in MS research.

In this work, we aim to elucidate the role of LPA_1_ in MS pathology, by analyzing the evolution of EAE disease course in maLPA_1_-null mice and in wild type mice in presence or absence of LPA_1_ antagonist. We provide evidence of a milder symptomatology in absence or antagonism of LPA_1_ receptor, suggesting a role of this receptor in the pathogenesis of the disease. This role was further strengthened with the analysis of LPA_1_ expression levels in peripheral blood mononuclear cells (PBMCs) from EAE mice and from patients with relapsing-remitting (RR-) MS, primary progressive (PP-) MS or secondary progressive (SP-) MS. We demonstrated that the initiation of relapses is accompanied by an increase of mouse LPA_1_ transcripts (*Lpar1*) in PBMCs. Finally, we provide *in vitro* data demonstrating that pro-inflammatory activation of human monocyte-derived macrophages includes increased expression of human LPA_1_ transcripts (*LPAR1)* and that LPA is involved in LPA_1_-driven polarization of human macrophages towards a M1-like phenotype. These results evidence a role of LPA in the initiation of the inflammatory process during MS relapses via LPA_1_.

In short, our studies unveil a novel mechanism for LPA in the classical activation of macrophages through LPA_1_, and suggest for the first time that targeting LPA_1_ receptors represents a promising therapeutic strategy in MS as well as for other immune-related diseases.

## Material and Methods

### Mice

Procedures were carried out with wild-type and maLPA_1_-null homozygous females (on a mixed background C57BL/6J × 129X1/SvJ) in compliance with European animal research laws (European Communities Council Directives 86/609/EU, 98/81/CEE, 2003/65/EC and Commission Recommendation 2007/526/EC) and national laws on laboratory animal welfare and approved by the local Experimentation Ethics Committees. The maLPA_1_-null (*Málaga* variant of LPA_1_-null; (12) mouse colony arose spontaneously from the initially reported LPA_1_-null mouse line (10) while crossing heterozygous foundational parents within their original mixed background. Experiments were performed on 7-week-old female mice obtained from heterozygous×heterozygous/homozygous maLPA_1_-null mating and genotyped for *Lpar1* deletion by PCR (10) or immunohistochemistry (12). Female maLPA_1_-null and wild-type mice were bred and housed in pathogen-free conditions at constant temperature (22 ± 1°C) and relative humidity (50%) under a regular light–dark schedule (light 7 am–7 pm). Food and water were freely available.

### Induction of EAE

Seven-week-old female mice were immunized according to a standard protocol (28) with subcutaneous injection of incomplete Freund’s adjuvant containing 4 mg/mL *Mycobacterium tuberculosis* (strain H37Ra; Difco Laboratories, Detroit, Michigan, USA) and 200 μg of encephalitogenic myelin oligodendrocyte glycoprotein peptide 35–55 (MOG_35–55_). The mice received intraperitoneal injections with 200 ng pertussis toxin (Sigma-Aldrich, St. Louis, Missouri, USA) at the time of immunization and 48 hours later. After 7 days, the mice received a half booster immunization with MOG/CFA and pertussis toxin. Control mice received identical injections without MOG_35–55_. Clinical disease usually commences around day 15 after immunization.

### LPA_1_ antagonist administration

VPC 32183 (S), (S)-Phosphoric acid mono-(2-octadec-9-enoylamino-3-[4-(pyridine-2-ylmethoxy)-phenyl]-propyl) ester (Ammonium Salt) (857340; Avanti Polar Lipids, Alabaster, Alabama, USA) was dissolved in 3% free-fatty acid BSA (FFA-BSA; Sigma-Aldrich) in saline. VPC32183 was diluted to a concentration of 5 µM and 100 µl were injected intravenous in the tail vein at the time-points described in the text. In non-treated control mice only vehicle injections were performed (3% FFA BSA in saline).

### Clinical evaluation

The mice were scored four times per week as follows: limp tail or waddling gait with tail tonicity, defined score 1; waddling gait with limp tail (ataxia) as score 2; ataxia with partial limb paralysis as score 2.5; full paralysis of one limb as score 3; full paralysis of one limb with partial paralysis of second limb as score 3.5. Animals that maintained a score of at least 3.5 more than 3 days were euthanized.

### Subjects

The samples for the RNA analysis of total PBMC were provided by the Biobank of our hospital, as part of the Andalusian Public Health System Biobank. All patients participating in the study gave their informed consent and protocols were approved by institutional ethical committees (Comite de Ética de la Investigación provincial de Málaga). The study was conducted according to international ethical principles contained in the Declaration of Helsinki (Fortaleza, 2013), Spanish regulations on biomedical research (Law 14/2007, of July 3, on biomedical research) and the provisions of the European General Personal Data Protection Regulation (Royal Decree-Law 5/2018, of 27 July, and Regulation (EU) 2016/679, of April 27, 2016). Patient selection was based on the criteria of first diagnosis and under no MS treatment.

For the RNA sequencing analysis, 22 MS patients (of which 32 were pairs of siblings) and 10 healthy controls were recruited for the macrophage experiments. The study was approved by the French Ethics committee and the French ministry of research (DC-2012-1535 and AC-2012-1536). Written informed consent was obtained from all study participants. All patients fulfilled diagnostic criteria for MS, and individuals (MS patients and healthy donors) with any other inflammatory or neurological disorders were excluded from the study.

### Isolation of PBMC for RNA extraction

PBMCs were isolated from whole blood by standard Ficoll®-Paque density gradient centrifugation. Briefly, heparinized blood was diluted with saline solution (1:1 dilution). Then, Ficoll®-Paque was covered with a layer of diluted blood. After 30 min of centrifugation (2000 rpm, room temperature (RT), without break), the PBMCs could easily be collected. After two washing steps and counting, cell were resuspended in 1 ml of Trizol® to extract RNA.

### Isolation of brain infiltrating mononuclear cells for RNA extraction

Infiltrated mononuclear cells (IMNCs) were isolated from CNS of EAE mice using the following procedure (6). After dissecting brain and spinal cord from individual animals, these were minced finely in phosphate buffer saline, centrifuged and resuspended in 37% Percoll®. This suspension was layed on a 70% Percoll® cushion and spun at 600 × g at room temperature for 25min. CNS IMNCs were obtained from the 37–70% Percoll® interface, washed twice, and cell counted. Finally, cells were resuspended in 1 ml of TRIzol® for RNA extraction.

### RT-PCR of PBMC and IMNC

Total RNA was extracted from PBMCs and IMNCs using the TRIzol™ reagent (Invitrogen Life Technologies) as originally described by Chomczynski and Sacchi (1987). cDNA was synthesized using 1 µg of total RNA by the enzyme reverse transcriptase MMLV (Sigma-Aldrich) and random primers. Real-time PCR was performed by the LightCycler® System (Roche Diagnostics) following the manufacturer’s specifications. The 10 µl final reaction volume consisted of 5.4 μl of distilled water RNAase-free, 1.3 μL of MgCl_2_, 0.2 μl of each forward and reverse primers, 1 μl of Fast SYBR™ Green Master Mix and 2 μl of cDNA. Reaction conditions were as follows: polymerase activation at 95°C for 15 min, 40 denaturation cycles of 95°C for 30 s, and annealing/elongation at 68°C (*Lpar1* and GAPDH) for 5 s (*Lpar1*) or 10 s (GAPDH).

The primer sequences used in the amplification of *Lpar1* and *Lpar2* have been described previously (Hama et al. 2004) (*Lpar1* forward: gaggaatcgggacaccatgat; and reverse: acatccagcaataacaagaccaatc, Gapdh forward: gccaaggtcatccatgacaact, and reverse: gaggggccatccacagtctt). A melting curve analysis was performed to assess primer specificity and product quality by step-wise denaturation of the PCR product at a rate of 0.1°C/sec to 98°C. The relative levels of receptor expression were quantified using the standard curve method.

### Isolation of Primary Monocytes and macrophage culture and activation

Blood was sampled from all participants in acid citrate dextrose (ACD) tubes. From blood samples, peripheral blood mononuclear cells (PBMCs) were isolated using Ficoll Paque Plus (GE Healthcare Life Sciences) and centrifugation (2200 rpm, 20 min without brake). Cells were washed in PBS (2×10 min at 1500 rpm) and RPMI 1640 + 10% fetal bovine serum (FBS) (5 min at 1500 rpm) (all products from ThermoFisher). Monocytes were isolated with anti-CD14 microbeads (Miltenyi) and plated in 12-well plates (500 000 cells/well) or in 24-well plates (200 000 cells/well) in RPMI 1640 + 10% FBS and granulocyte macrophage colony-stimulating factor (GM-CSF) (500 U/ml, ImmunoTools). After 72h, media was replaced with fresh media and one of the following: GM-CSF (500 U/ml); IFNβ (100 U/ml, ImmunoTools); IL-4 (1000 U/ml, ImmunoTools); or combined IFNγ (200 U/ml, ImmunoTools) and ultra-pure LPS (10 ng/ml, InvivoGen). Cell lysis and RNA extraction were performed 24h post-activation using Nucleospin RNA extraction kit (Macherey-Nagel). Quality of RNA was confirmed on Agilent TapeStation (RINe>8).

For LPA treatment and antagonist, LPA (Tocris, 3854) and Ki16425 (Sigma, SML0971) were dissolved in 3% BSA to add to the medium at a final concentration of 1µM LPA and 400nM Ki16425 during 24h.

### RNA sequencing

Transcriptome sequencing cDNA libraries of macrophage RNA were prepared using a stranded mRNA polyA selection (Truseq stranded mRNA kit, Illumina). For each sample, we performed 60 million single-end, 75 base reads on a NextSeq 500 sequencer (Illumina). RNA-Seq data analyses were performed by GenoSplice technology (www.genosplice.com). Sequencing, data quality, reads repartition (e.g., for potential ribosomal contamination), and insert size estimation were performed using FastQC, Picard-Tools, Samtools and rseqc. Reads were mapped using STARv2.4.0 (11) on the hg19 Human genome assembly. Gene expression regulation study was performed as previously described (34). Briefly, for each gene present in the FAST DB v2018_1 annotations, reads aligning on constitutive regions (that are not prone to alternative splicing) were counted. Based on these read counts, normalization was performed using DESeq2 (26) in R (v.3.2.5). Genes are considered as expressed if their RPKM value is greater than 97.5% of the background RPKM value based on intergenic regions. The normalized data were used for all subsequent analysis.

### RT-PCR of monocyte-derived macrophages

RNA obtained from differentially activated macrophages were used as templates to synthetize cDNA using the Quantitect Reverse Transcription kit (cat No./ID: 205313) according to the manufacturer’s protocol. Quantitative RT-PCR was performed with the LightCycler® 1536 Instrument (Roche), and the following primers:

Hs_CD86_1_SG QuantiTect Primer Assay (200) Cat No./ID: QT00033915

Hs_TLR2_1_SG QuantiTect Primer Assay (200) Cat No./ID: QT00236131

Hs_CCL2_1_SG QuantiTect Primer Assay (200) Cat No./ID: QT00212730

Hs_CCL5_1_SG QuantiTect Primer Assay (200) Cat No./ID: QT00090083

Hs_CCL20_1_SG QuantiTect Primer Assay (200) Cat No./ID: QT00012971

Hs_*LPAR1*_1_SG QuantiTect Primer Assay (200) Cat No./ID: QT00021469

Hs_MRC1_1_SG QuantiTect Primer Assay (200) Cat No./ID: QT00012810

Hs_CD163_1_SG QuantiTect Primer Assay (200) Cat No./ID: QT00074641

Hs_CD180_1_SG QuantiTect Primer Assay (200) Cat No./ID: QT00203574

Hs_PDGFC_1_SG QuantiTect Primer Assay (200) Cat No./ID: QT00026551

### Statistical methods

All mouse studies were repeated a minimum of 3 times, and each experimental group included at least 4 samples. The values are expressed as mean ± SEM. The statistical analysis was done with Graphpad. Statistical significance was determined using the appropriate statistical test mentioned in each experiment. Values were considered to be statistically significant when p< 0.05.

## Results

### LPA_1_ deletion leads to milder EAE clinical course

Recently, a role of LPA in the pathogenesis of the MS and its animal model, EAE, has been suggested (41), focusing on the contribution of LPA_2_-expressing T cells. Here, we question whether LPA_1_, another LPA receptor, also present in immune cells, could have a role in EAE. To answer this question, we first compared the EAE clinical course in presence and absence of LPA_1_ by comparing MOG_35-55_ induced-EAE in wild type and in the *Malaga* variant of LPA_1_-null mouse line (maLPA_1_-null mouse) (12).

Analysis of their clinical courses showed a relapsing-remitting clinical course in both wild-type and LPA_1_-null animals but also highlighted also important differences between the two genotypes. Notably, maLPA_1_-null mice showed a less severe clinical course compared to wild-type mice (Fig. 1A), measured as the area under the curve (AUC). In addition, maLPA_1_-null mice exhibited a significantly lower average clinical score and maximal clinical score reached during relapses as well as a better recovery during remission (Fig.1B).

**Fig. 1.**
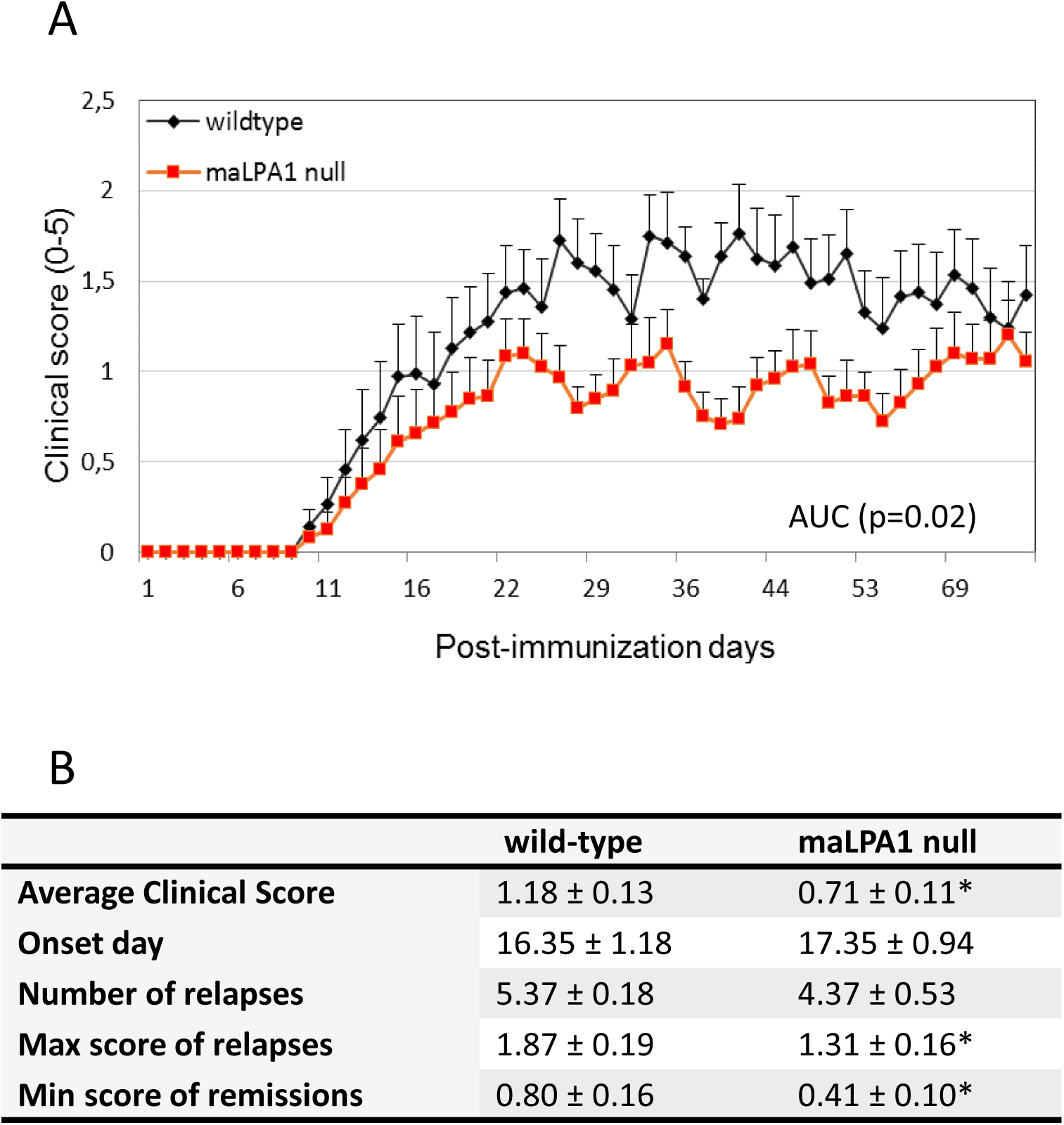
LPA_1_ null mice exhibit a less severe EAE disease course than wild type (WT). A) EAE disease progression in wild-type (n=14) and maLPA_1_ knockout (n=20) mice. Graphs present mean values with error bars indicating + SEM of summarizes data from three independent experiments. Statistical significance was assessed by Mann-Whitney test of AUC (p=0.027) B) Comparison of clinical parameters of EAE in wild-type and maLPA_1_ null mice. T-student test for the average clinical score p=0.01; for the max score p=0.03; and for the min score p=0.04. * means p<0.05.

### Intravenous injection of an LPA_1_ antagonist ameliorates EAE clinical score in wild-type mice

LPA_1_ is also expressed in oligodendrocytes and we previously demonstrated that its absence perturbs developmental myelination in maLPA_1_-null mice (14). To exclude any interference of deficient myelin patterns in the EAE outcome in LPA_1_-null animals, we used a pharmacological approach in wild-type mice, intravenously injecting an LPA_1_ antagonist (VPC32183) that primarily blocks LPA_1_ (10-100nM range), and partially blocks LPA_3_ (10-fold lower; 100-1000nM range) (17).

We first administered a single dose of the antagonist at the clinical onset of the disease, at 14 days post-immunization (dpi). This protocol resulted in a trend towards an amelioration during the first 5 days (Fig. 2A). However, after that period, the symptoms reappeared more severely. Due to its lipidic structure, LPA_1_ antagonist could easily be metabolized in the blood stream, explaining the short duration of its positive effects.

**Fig. 2.**
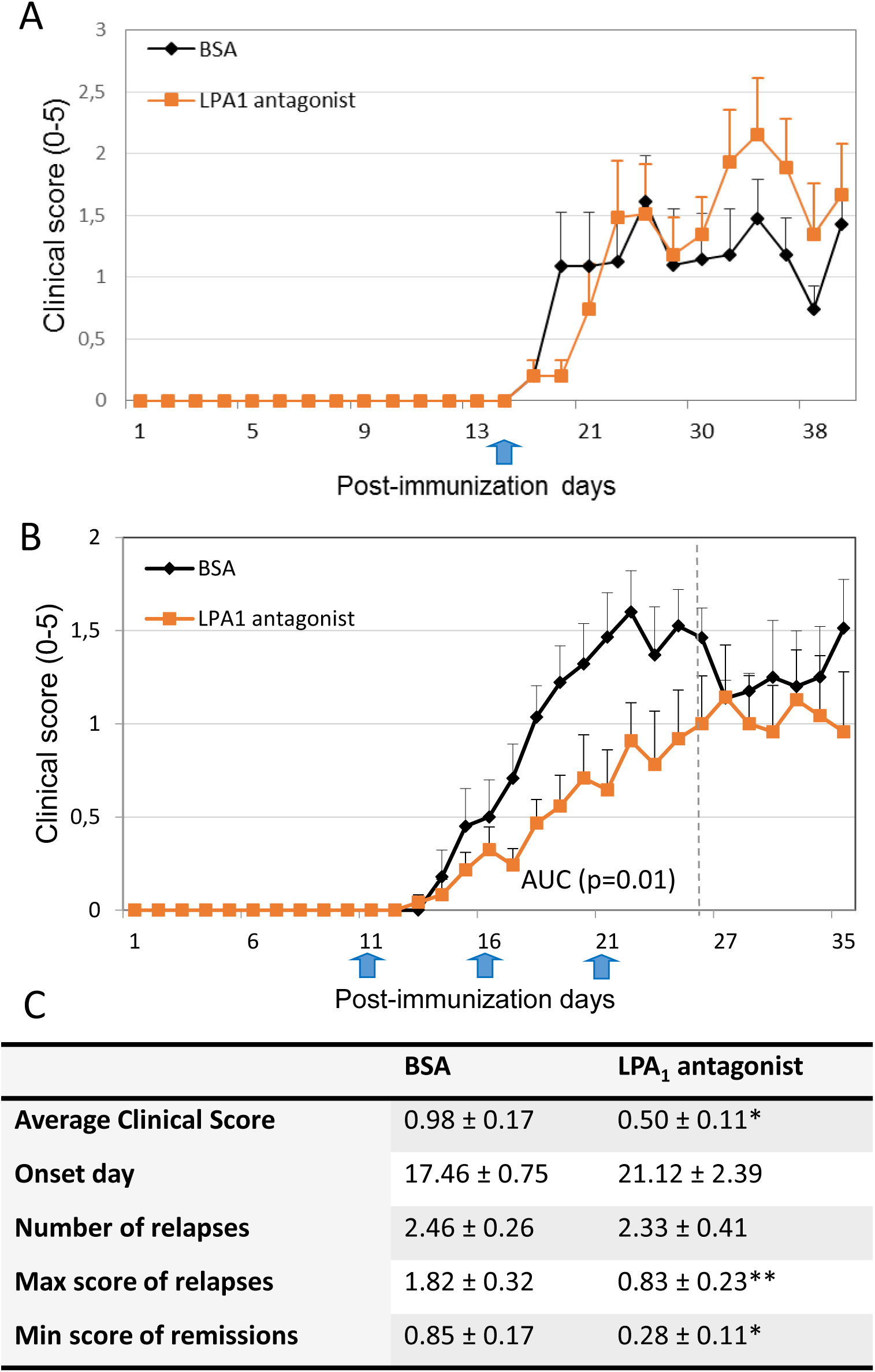
LPA_1_ antagonist treatment lowers clinical scores in EAE mice. A) EAE disease progression in wild-type mice with one LPA_1_ antagonist dose at 14dpi (blue arrow) (BSA group n=8, LPA_1_ antagonist group n=4). B) EAE disease progression in wild-type mice with three LPA_1_ antagonist doses at 11, 16 and 21dpi (blue arrows). The graph summarizes data from three independent experiments with total n of 14 and 12 of BSA and LPA_1_ antagonist-treated mice respectively. Graphs present mean values with error bars indicating + SEM. Statistical significance was assessed by Mann-Whitney test of AUC up to 25dpi (p=0.019) C) Comparison of clinical parameters of EAE in vehicle- and LPA_1_ antagonist – treated wildtype animals Mann Whitney test: average clinical score p=0.046, max score of relapses p=0.001, min score of remissions p=0.032. * means p<0.05, ** means p<0.01.

To maintain the levels of the antagonist, new sets of immunized mice were treated with repeated doses of VPC32183 every 5 days (Fig. 2B). Recurrent intravenous administration of the LPA_1_ antagonist resulted in a milder EAE disease course characterized by a lower average clinical score, milder relapses and better remissions (Fig. 2B,C), corroborating the requirement of LPA_1_ activation to develop a normal EAE clinical course. Nevertheless, after ceasing the treatment (25dpi), the severity of the clinical course went back to control levels (Fig.2B), indicating a reversible effect of the antagonist.

### *Lpar1* expression increases when mononuclear cells invade the CNS

Under normal physiological conditions, mononuclear cells are rarely found in the CNS. However, in MS and EAE, activated immune cells infiltrate the CNS. Due to the reported role of LPA_1_ in immune cells infiltration (42), and its obvious impact in the clinical course, we analyzed whether the number of infiltrating mononuclear cells (IMNC) was altered in EAE-mice lacking LPA_1_. We quantified the number of IMNCs isolated by Percoll gradient (6) from brain and spinal cord of wild type and maLPA_1_-null mice with similar clinical score. We did not find significant differences in the number of infiltrates (Fig. 3A), suggesting that LPA_1_ modulation of EAE course might intervene at another step of immune cell activation beside infiltration.

**Fig. 3.**
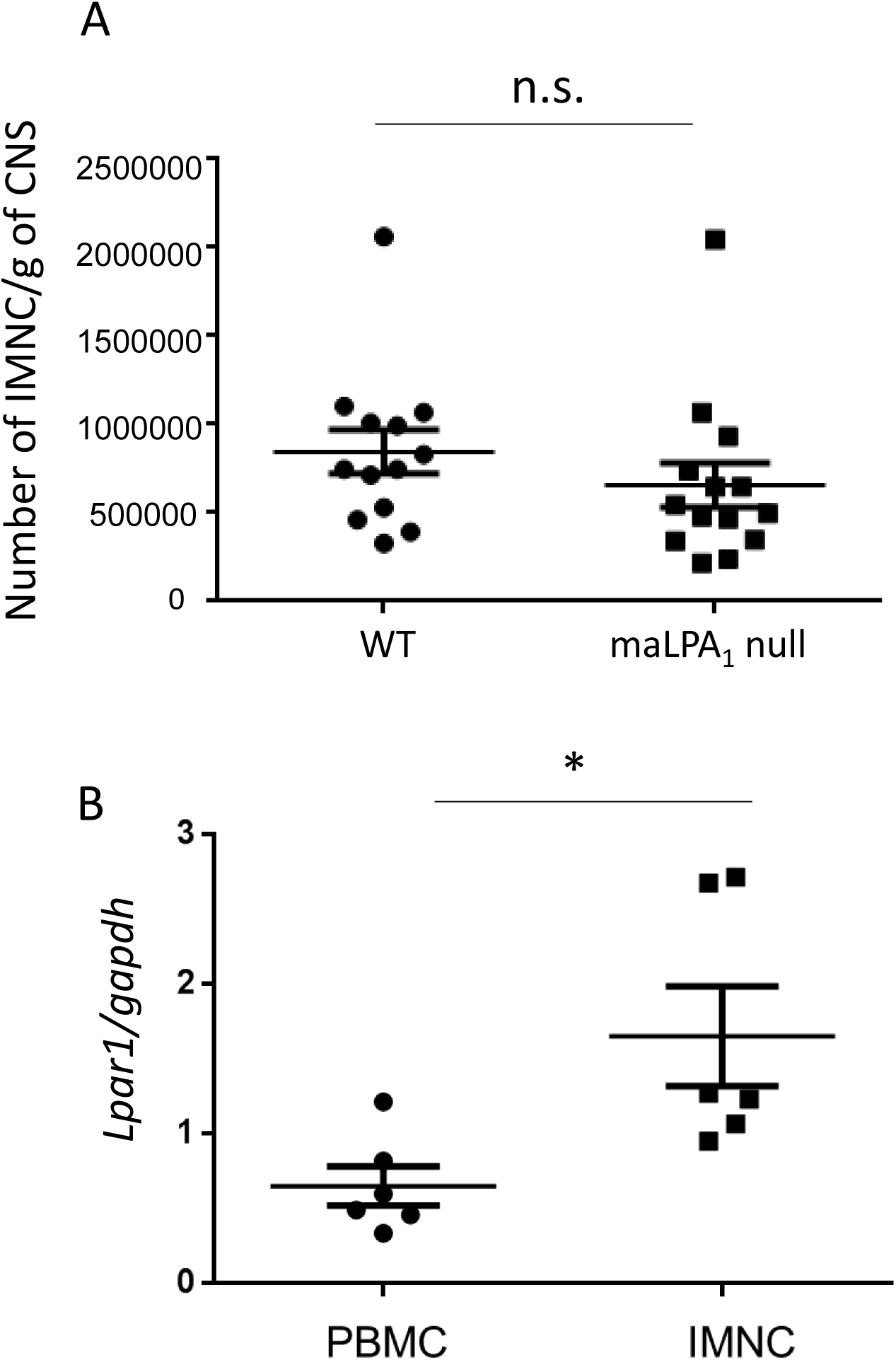
*Lpar1* increases in CNS infiltrating macrophages compared to circulating PBMC but does not participate to macrophage recruitment. A) There was no difference in the number of IMNC within brain and spinal cord of two sets of wt and maLPA_1_ ko mice with similar clinical courses. Clinical course average of wildtype mice (n=13) was 1.36±0.19 and maLPA_1_ (n=14) was 1.43±0.17. IMNC was normalized by CNS weight (g): wt=845276±124261; maLPA1=658148±125219. Unpair t-test, p=0.29. B) *Lpar1* expression normalized to gapdh was higher in CNS IMNC when compared to the *Lpar1* levels in PBMC (n=6 per group) Clinical course average was 1.56±0.4. Wilcoxon matched-pair test p=0.031. * means p< 0.05, n.s. means not significant.

Despite LPA_1_ not being essential for infiltration, its expression still appeared to be related to the EAE pathogenesis. Thus, we analyzed the *Lpar1* expression in circulating immune cells (PBMCs) and CNS-IMNCs from wild-type mice using RT-PCR and our results show an increase of *Lpar1* expression after immune cell infiltration in the CNS compared to PBMCs (Fig. 3B). These data suggest that an increase of *Lpar1* expression reflects immune cell activation.

### Onset of EAE relapses correlates with increase expression of *Lpar1* in PBMCs

Knowing that *Lpar1* is expressed by immune cells, and that immune cells are critical for EAE development, we wondered whether *Lpar1* expression in PBMCs reflects disease activity. To this end, the expression of *Lpar1* in PBMCs along the EAE clinical course was evaluated in wild-type mice.

MOG-immunized animals showed a two-fold significant increase of *Lpar1* expression compared to control animals (Fig. 4A). However, no significant correlation was found between *Lpar1* expression and clinical score after sacrifice (EAE score 1: 0,459818 ± 0,123361 (n=9); EAE score 2: 0,681039 ± 0,151682 (n=7); EAE score 3: 0,648933 ± 0,144792 (n=8)).

**Fig. 4.**
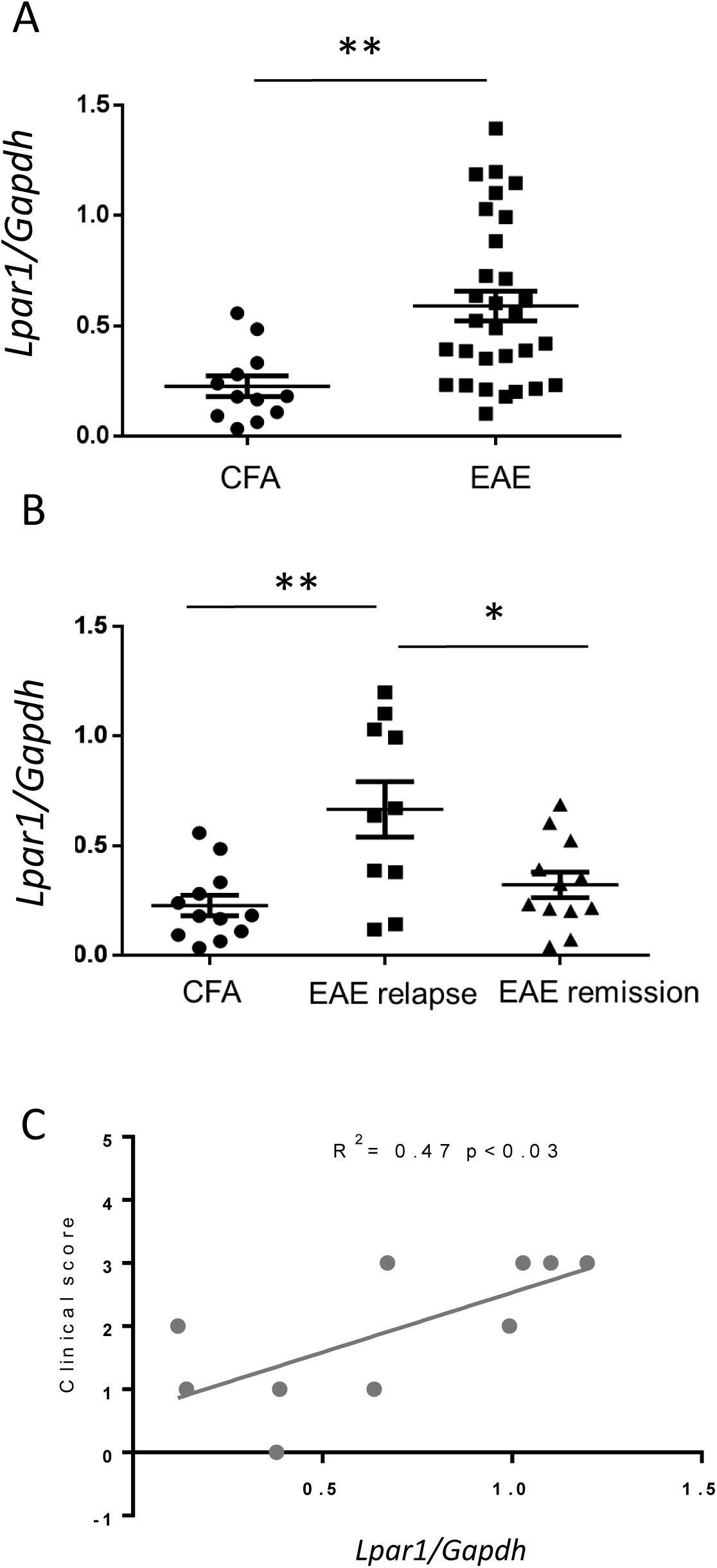
*Lpar1* expression in mouse PBMC during EAE development. A) Relative RT-PCR analysis of *Lpar1* expression in PBMC normalized to *Gapdh* in control (n=12) and EAE-induced (n=30) animals regardless the moment of the disease. The expressions of *Lpar1* in EAE-induced mice were double to those in control animals. t-student p= 0.0022. B) The *Lpar1* expression of the EAE mice sacrificed during relapses (n=10) was significantly higher than in controls (t-student test p=0.0024) while *Lpar1* expression reduced during remission (n=12) (t-student test between relapses and remissions groups p=0.016). C) Positive correlation between the clinical symptoms and the expression of *Lpar1* in EAE mice during relapses.

To decipher whether *Lpar1* expression in PBMCs might reflect a different phase of the disease, *Lpar1* expression was analyzed according to mice stratification based on whether animals were initiating a relapse or in remission/progressive course of the disease at the moment of the sampling. There was a significantly increased expression of *Lpar1* during relapses when compared to control animals or animals in remission or progressive episodes (Fig. 4B). Of note, *Lpar1* expression and clinical symptoms of EAE were significantly positively correlated during the clinical course of the disease (Fig. 4C).

### *LPAR1* expression increases during relapses in RRMS patient PBMCs

Our previous observations in EAE mice suggested a modulation of LPA_1_ in the first stages of the relapses during the inflammatory clinical course. To corroborate this observation in the context of MS, we compared the expression of *LPAR1* in PBMCs from RR-MS patients at the time of first relapse and compared with its expression in healthy donors (HD), matched in age and gender (Fig. 5A), and patients with progressive form of the disease (SP-MS and PP-MS). Like in EAE, *LPAR1* expression was significantly higher in RR-MS patient PBMCs than in healthy subjects or progressive patients (Fig. 5B). Thus, we provide evidence that alterations in *LPAR1* expression associates with the inflammatory phase of MS.

**Fig. 5.**
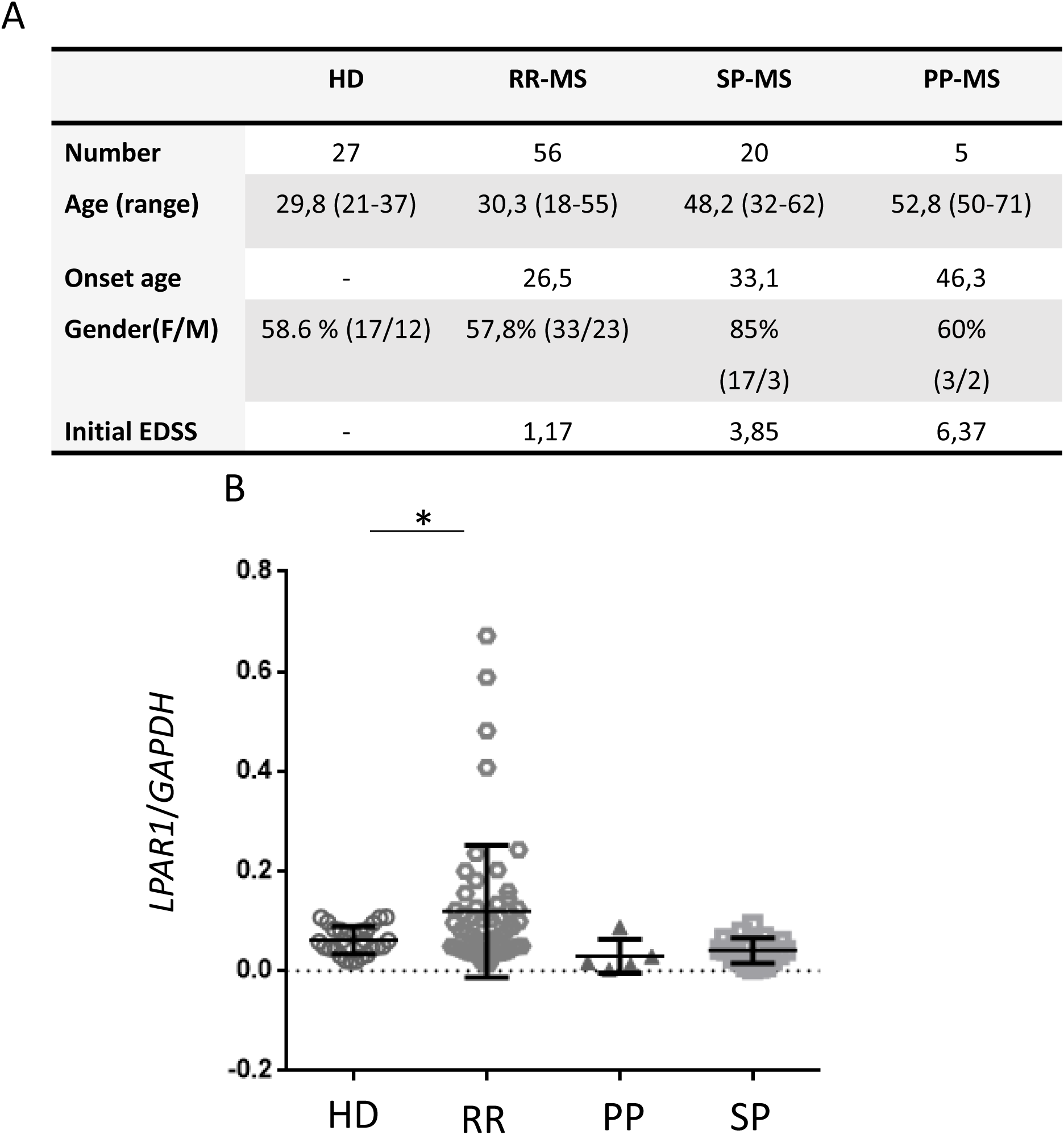
Up regulation of LPA_1_ expression in MS patient PBMCs during relapses. A) Demographic data of the studied groups. B) Relative expression of *LPAR1* normalized to *GAPDH* in healthy donor, RR-MS patients during relapses, SP-MS and PP-MS patients. One-way ANOVA (p=0.004, Bonaferri posthoc test)

### *LPAR1* expression correlates with a pro-inflammatory phenotype of human monocyte-derived macrophages

Circulating PBMCs are mainly composed of lymphocytes and monocytes. We decided to focus on monocytes/macrophages because of their dual role in MS pathology (46), being both deleterious when endorsing a pro-inflammatory phenotype and beneficial under pro-regenerative activation (30, 31).

To elucidate the role of LPA_1_ in macrophages polarization, we obtained naïve circulating monocytes from RR-MS patients in remission. Naïve monocytes remain circulating in the blood stream for a short period before infiltrating the tissues (38), reducing the impact of other circulating factors before blood extraction. Blood CD14+ monocytes were differentiated into macrophages *in vitro* using GMCSF before testing the role of LPA in macrophage polarization.

In order to elucidate how LPA_1_ might correlate the macrophage activation state, monocyte-derived macrophages from healthy controls (Fig 6, circles) or MS patients (Fig 6, triangle) were directed toward a pro inflammatory (LPS+IFNγ, Fig 6, blue), a pro regenerative state (IFNβ (Fig 6, purple) or IL-4 (Fig 6, red)) or a neutral state (GMCSF, Fig 6 green). We next evaluated by RNA sequencing, the expression of three LPA receptors: two membrane receptors, LPA_1_ and LPA_2_, and a nuclear receptor PPARγ.

**Fig. 6.**
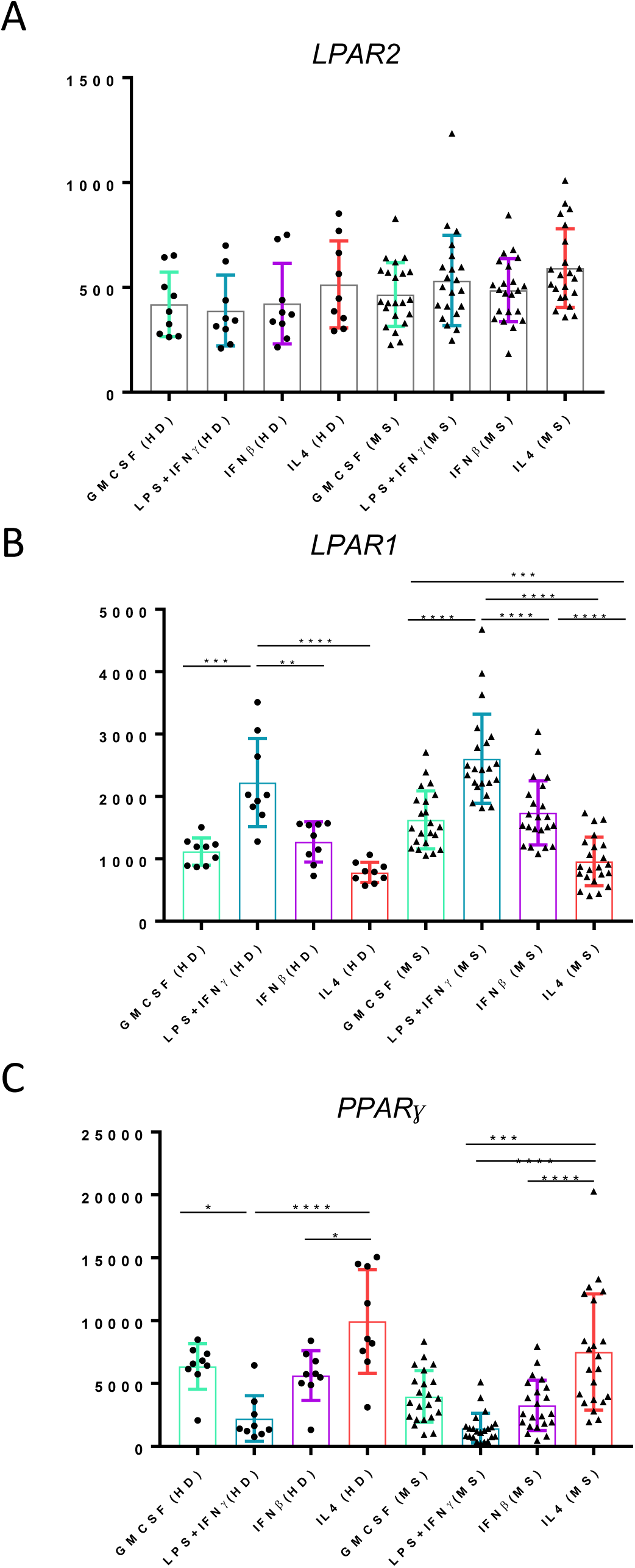
LPA receptor LPA_1_ and PPARγ are differentially regulated in human macrophages after pro inflammatory and pro regenerative differentiation. RNA sequencing analysis. Comparison of macrophage expression profiles in naïve (GMCSF), classically activated (LPS+INFγ) or alternatively (IFNβ or IL4) human macrophages, from HD (n=9) and RR-MS patients (n=22). While *LPAR2 expression* did not change after any type of activation (A), *LPAR1* expression was significantly increased in both HD and RR-MS patients after pro inflammatory activation (B). In contrast, the nuclear LPA receptor *PPARγ* was increased in the pro-regenerative state and significantly reduced in pro inflammatory macrophages in HD but not in RR-MS patients (C).

Interestingly, these receptors were differentially regulated, while we could not detect a difference of *LPAR2* expression (Fig. 6A) in any activation state, we observed an inverse pattern of expression for *LPAR1* (Fig 6B) and *PPARγ* (Fig. 6C) across the macrophage activation states. In both MS and HD, *LPAR1* expression is up-regulated in the pro-inflammatory state, while PPARγ expression is increased in the pro-regenerative state. This observation underlines the possible role of LPA_1_ in the pro-inflammatory activation of human macrophages.

### LPA mediates human macrophage polarization

In addition to the increase of LPA_1_ in pro-inflammatory macrophages (Fig. 6B), LPA levels are altered along the course of MS (4, 41) suggesting an important role of this phospholipid in the course of the disease. We therefore tested whether LPA could promote an M1-phenotype in macrophages, as has been observed in murine microglia (39).

We examined transcripts specific for pro-inflammatory or pro-regenerative profiles to test the effect of LPA treatment for 24h, and compared the expression of different markers with the canonical M1 polarization by LPS (Fig. 7).

**Fig. 7.**
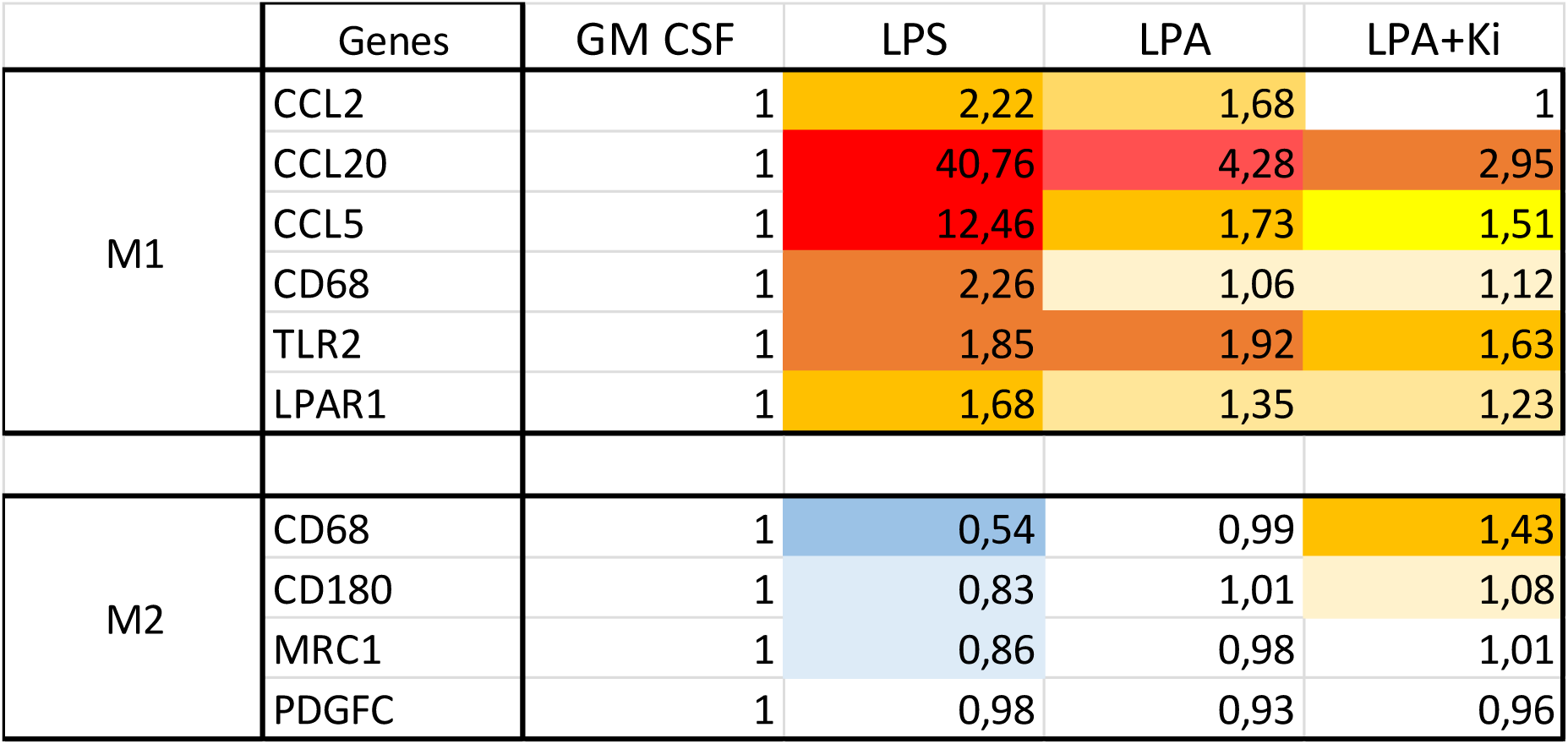
LPA_1_ antagonist directs human macrophages toward a more pro regenerative phenotype. Heatmap representing the expression of specific markers of pro-inflammatory or pro-regenerative phenotypes after macrophages activation with LPS, LPA and LPA+Ki16425, expressed as ratio to the non-activated condition (GMCSF). LPA treatment (1µM) increased the expression of the M1-like marker genes in a milder manner as compared to LPS. Addition of Ki16425 (400nM) reduced the M1-like polarization effect of LPA, indicating that this response is mediated (at least partially) by LPA_1_. Data are normalized to the housekeeping gene HPRT and represent as mean of three different individuals.

The levels of different M1 markers (CCL2, CCL20, CCL5, CD68 and TLR2), though to a lesser extent after LPS treatment, increased after LPA incubation, indicating a role of LPA in the pro-inflammatory activation of human macrophages. Moreover, this polarization was partially inhibited by addition of an LPA_1_ inhibitor (Ki16425) revealing the mediation of LPA_1_ in this LPA response (Fig. 7). No significant alterations in the expression of M2 markers were observed after LPA incubation.

## Discussion

In this study, we present evidence of a role of a receptor of LPA (a signaling molecule with a broad effector profile (9) and described roles in inflammation (48)) in the pathogenesis of the neuro-inflammatory disease MS and its animal model, EAE. We also propose a mechanism through which LPA may exert this effect via macrophage activation.

After the discovery of the first receptor for LPA, the LPA_1_, in 1996 (18), this receptor has been implicated in a numerous process, with an outstanding importance in the physiology and pathology of the CNS (49). In this context, the importance of LPA in MS pathogenesis, one of the broadest spread neuro-inflammatory diseases has been suggested (4, 21, 41). However, the role of LPA_1_ in the MS pathogenesis remains unclear. In the present study, we unveil a new aspect of LPA through the LPA_1_ receptor in this neuro-inflammatory disease.

Our results show for the first time, the importance of the receptor LPA_1_ in EAE clinical course. The lack of LPA_1_, or its pharmacological inhibition by the repetitive intravenous injections of a LPA_1_ antagonist (VPC32183), reduces the severity of the disease as seen by a lower average clinical score and lower maximal score during relapses as well as better recovery during remission. The milder symptoms in the absence of LPA_1_ signaling indicates that this pathway is involved in EAE pathogenesis. This is in contrast with a study of another LPA receptor, LPA_2_, of which reduction led to more severe disease (41). This indicates a complex role of LPA in MS and EAE, and that potential treatment strategies should target specific pathways rather than LPA as a whole.

Previous studies have described a role for LPA and autotaxin, its main synthetic enzyme, in inflammatory processes (8, 25, 43, 45). In line with this, we found that expression of LPA_1_ was high during relapses – which are generally associated with high inflammatory activity – in immune cells in both EAE and MS. The differential expression of *LPAR1* in the different clinical forms of MS indicates a direct role of the LPA-LPA_1_ pathway in the inflammatory component of the disease. In addition, this suggests a potential use of *LPAR1* expression as a biomarker of disease activity. These results mirror studies showing increased levels of LPA in blood or cerebrospinal fluid of RR-MS patients during relapse compared to healthy controls or RR-MS patients in remission (4, 21), and suggest a broader dysregulation of LPA signaling than previously thought.

We also found a significant positive correlation between the levels of *Lpar1* expression during relapses and the severity of the EAE clinical course, encouraging future analysis of RR-MS patient clinical disability and *LPAR1* expression. A correlation between the two would strengthen the implication of LPA_1_ in the disease course and potentially enable the use of its expression to estimate the individual patient’s prognosis. Nevertheless, a large cohort and consideration of any immune-modulatory treatment would be necessary to extract meaningful statements.

To understand how LPA_1_ exerts its influence on the inflammatory component of MS and EAE, we examined infiltration and activation of immune cells. In the case of LPA_2_, its effect on the EAE disease course appears to be reliant on its capacity to increase T-cell homing, thus reducing infiltration. While studies have indicated a detrimental role of LPA_1_ in blood-brain-barrier (BBB) integrity (5, 36, 47) and potential to increase extravasation through induction of chemokine expression (24), our results did not indicate a significant impact of LPA_1_ deletion on infiltration of PBMCs into the CNS. While this does not exclude an effect below statistical significance or an effect of BBB leakage independent of PBMC infiltration, we cannot explain the amelioration of clinical scores through reduced infiltration. Instead, the observation that infiltrating cells express *Lpar1* to a higher degree than peripheral cells in EAE wild-type mice suggests that LPA_1_ is involved in immune cell activation without necessarily affecting infiltration.

Following the hypothesis that LPA_1_ correlates with immune cell activation, we examined its expression in activated human macrophages. Our results show an increase of *LPAR1* expression, but not *LPAR2*, in both healthy control and RR-MS patient macrophages when activated towards a pro-inflammatory phenotype. On the other hand, expression of the LPA nuclear receptor *PPARγ* not only decreases when macrophages acquire M1 polarization but also shows a trend towards increasing after pro-regenerative activation. We did not identify a difference between MS patients and healthy controls, but this could be due to the fact that MS samples were taken during a remission phase. In this case, the modular expression of different LPA receptors after differential activation hints a complex role of LPA signaling in the homeostasis of macrophages during the disease and suggests that modulating the expression or saturating the activation of one or the other one could be a mechanism of trans-differentiation of human macrophages. Although this aspect requires further study, the fact that LPA_1_ is related to glycolysis (7, 16, 33) (main source of energy for pro-inflammatory polarized macrophages) and PPAR*γ* induces oxidative phosphorylation (35), and that these two metabolic processes are central in pro- and anti-inflammatory macrophage activation respectively (13), suggests that the modulation of these LPA receptors could have major implications in the macrophage physiology and activation.

Knowing that LPA is dysregulated in MS relapses (4, 21) and that *LPAR1* expression is increased in macrophage activation, we hypothesized that LPA_1_ could mediate LPA-induced pro-inflammatory activation in macrophages. This was confirmed through increased expression of M1 markers following LPA incubation with partial correction by exposure to the LPA_1_ antagonist Ki16425. Increased expression of LPA_1_ in EAE and MS PBMCs during relapse thus suggests both an activated state as well as a predisposition to further pro-inflammatory activation. The coordinated responses between the induced-LPA_1_ expression and the pro-inflammatory activation of LPA via the LPA_1_ will promote a positive feedback loop that grants to LPA the role of boosting the inflammatory response and maintain the classical activation of macrophages. The milder EAE clinical course observed in LPA_1_-null and LPA_1_-antagonized mice, which present lower maximal and minimal scores in relapses and remissions respectively, could therefore be explained with a milder activation of immune cells, whereas the number or relapses and the onset of the disease unaffected as infiltration still occurs to the same extent.

In short, our study unveils for the first time a role of the LPA_1_ in the pathogenesis of MS and its animal model, EAE, and the importance of the regulation of the LPA signaling in the development of the disease. In addition to opening up new avenues for immuno-modulatory treatment, this research also indicates a potential for LPA_1_ as a biomarker of disease activity. Further research on LPA in MS should therefore consider the exact pathways being targeted and the current level of disease activity in the patient, in order to develop strategies to better follow and treat these neurological patients.

## Acknowledgement

This work was supported by grants from Spanish Ministry of Science, Innovation and Universities, co-funded by European Regional Development Fund (ERDF, EU), (PI16/01510, to GET), Andalusian Regional Ministries of Economy, Knowledge, Business and University (Talent Hub Programme, co-funded by FP7,UE, to BGD; CTS643 to GET) and of Health and Families (Nicolas Monardes Programme, and PI-0187/2008, PI-0234-2013 to GET), and Ramon Areces Foundation to BGD. This work was supported by the OCIRP foundation, the program “Investissements d’Avenir” ANR-10-IAIHU-06 and “Translational Research Infrastructure for Biotherapies in Neurosciences” ANR-11-INBS-0011–NeurATRIS.

We also gratefully acknowledge IBIMA joint support structures for research (ECAI) of General Services, Microscopy and Animal Experimentation. For the ICM, we would like to acknowledge the center for clinical investigation, the IGENSEQ sequencing platform and the CELLIS cell culture facility but also the fundraising, scientific affairs and the administrative department.

